# Highly Multiplexed Single-Cell Full-Length cDNA Sequencing of human immune cells with 10X Genomics and R2C2

**DOI:** 10.1101/2020.01.10.902361

**Authors:** Roger Volden, Christopher Vollmers

## Abstract

Single cell transcriptome analysis elucidates facets of cell biology that have been previously out of reach. However, the high-throughput analysis of thousands of single cell transcriptomes has been limited by sample preparation and sequencing technology. High-throughput single cell analysis today is facilitated by protocols like the 10X Genomics platform or Drop-Seq which generate cDNA pools in which the origin of a transcript is encoded at its 5’ or 3’ end. These cDNA pools are most often analyzed by short read Illumina sequencing which can identify the cellular origin of a transcript and what gene it was transcribed from. However, these methods fail to retrieve isoform information. In principle, cDNA pools prepared using these approaches can be analyzed with Pacific Biosciences and Oxford Nanopore long-read sequencers to retrieve isoform information but current implementations rely heavily on Illumina short-reads for analysis in addition to long reads. Here, we used R2C2 to sequence and demultiplex 12 million full-length cDNA molecules generated by the 10X Chromium platform from ∼3000 peripheral blood mononuclear cells (PBMCs). We used these reads to – independent from Illumina data – cluster cells into B cells, T cells, and Monocytes and generate isoform-level transcriptomes for these cell types. We also generated isoform-level transcriptomes for all single cells and used this information to identify a wide range of isoform diversity between genes. Finally, we also designed a computational workflow to extract paired adaptive immune receptors – T cell receptor and B cell receptor (TCR and BCR) – sequences unique to each T and B cell. This work represents a new, simple, and powerful approach that – using a single sequencing method – can extract an unprecedented amount of information from thousands of single cells.

## Introduction

The analysis of transcriptomes using high-throughput sequencers has revolutionized biomedical research [1,2]. Pairing transcriptome analysis with the high-throughput processing of single cells has provided unprecedented insight into cellular heterogeneity [3,4]. Among many other studies, researchers have leveraged the strengths of high-throughput single-cell transcriptome analysis to create single cell maps of the mouse [5,6] or C. elegans [7] model organisms, to elucidate a new cell type in the lung involved in cystic fibrosis [8], and to increase our knowledge of adaptive and innate immune cells [9–12].

High-throughput single-cell transcriptome analysis however comes with trade-offs. In particular, droplet- or microwell-based methods like Drop-seq [13], 10X Genomics [14], and Microwell-Seq [6] or Seq-Well [15] single cell workflows generate pools of full-length cDNA with either the 5’ or 3’ end containing cellular identifiers. The cDNA pools are intended for high-throughput short-read sequencing and must therefore be fragmented such that one read sequence includes the cellular identifier and the sequence of its pair includes a fragment from within the original cDNA molecule. As a result, only a relatively short fragment of the cDNA is then sequenced alongside the cellular identifier limiting the resolution of this approach to the identification of genes associated with a given molecular identifier.

Instead of sequencing transcript fragments, long-read sequencing methods in the form of Pacific Biosciences (PacBio) and Oxford Nanopore Technologies (ONT) are now capable of sequencing comprehensive full-length transcriptomes [16–19]. These methods have now been used to analyze single cell cDNA pools generated by different methods, both well- [20–22] and droplet-based [23–25], enriching the information we can extract from single cells experiments. However, for the analysis of high-throughput droplet-based experiments with long reads, short-read data are still required for interpreting experimental data [26] or enabling the identification of cellular and molecular identifiers in low-accuracy ONT reads [24]. Short-read data remain a requirement because either long-read data are not of sufficient depth to cluster cells into cell types or not accurate enough to decode cellular origin of cDNA molecules.

Because decoding the cellular origin of a cDNA molecule requires accurate sequencing of the molecular identifier, error-prone long read technologies are generally not sufficient to sequence each cDNA pool and to accurately interpret the single-cell data encoded therein. We have recently developed and applied the R2C2 approach which uses concatemeric consensus sequencing to improve ONT read accuracy from ∼92% to >99% while still producing more than 2 million full-length cDNA sequences per MinION flow cell [19,20,27,28]. The combination of these technologies therefore has the potential to illuminate isoform-level single cell biology with unprecedented resolution.

In this manuscript we demonstrate that this combination of high throughput and accuracy is sufficient for the Illumina short-read independent analysis of highly multiplexed 10X Genomics cDNA pools. To this end we independently analyzed two pools containing the cDNA molecules a combined ∼3000 human Peripheral Blood Mononuclear Cells (PBMCs) with Illumina and R2C2 (ONT) workflows. We showed that the R2C2 approach identifies the same cellular identifiers in the cDNA pools and generates comparable single-cell gene expression profiles and cell type clusters. In addition, and in contrast to Illumina data, R2C2 data also allow the determination of cell type specific and single-cell isoform-level transcriptomes. Finally, R2C2 allowed us to resolve and pair full-length adaptive immune receptors (AIR) transcripts in the B and T cell subpopulations of our PBMC sample which currently requires specialized library preparation methods and sequencing approaches.

## Results

We extracted PBMCs from whole blood and processed the cells in replicate using the Chromium Single Cell 3’ Gene Expression Solution (10X Genomics) aiming to include 1500 cells each for two replicates. We then divided the full-length cDNA intermediate generated by the standard 10X Genomics protocol to perform both short- and long-read sequencing (Figure 1A).

**Fig. 1:**
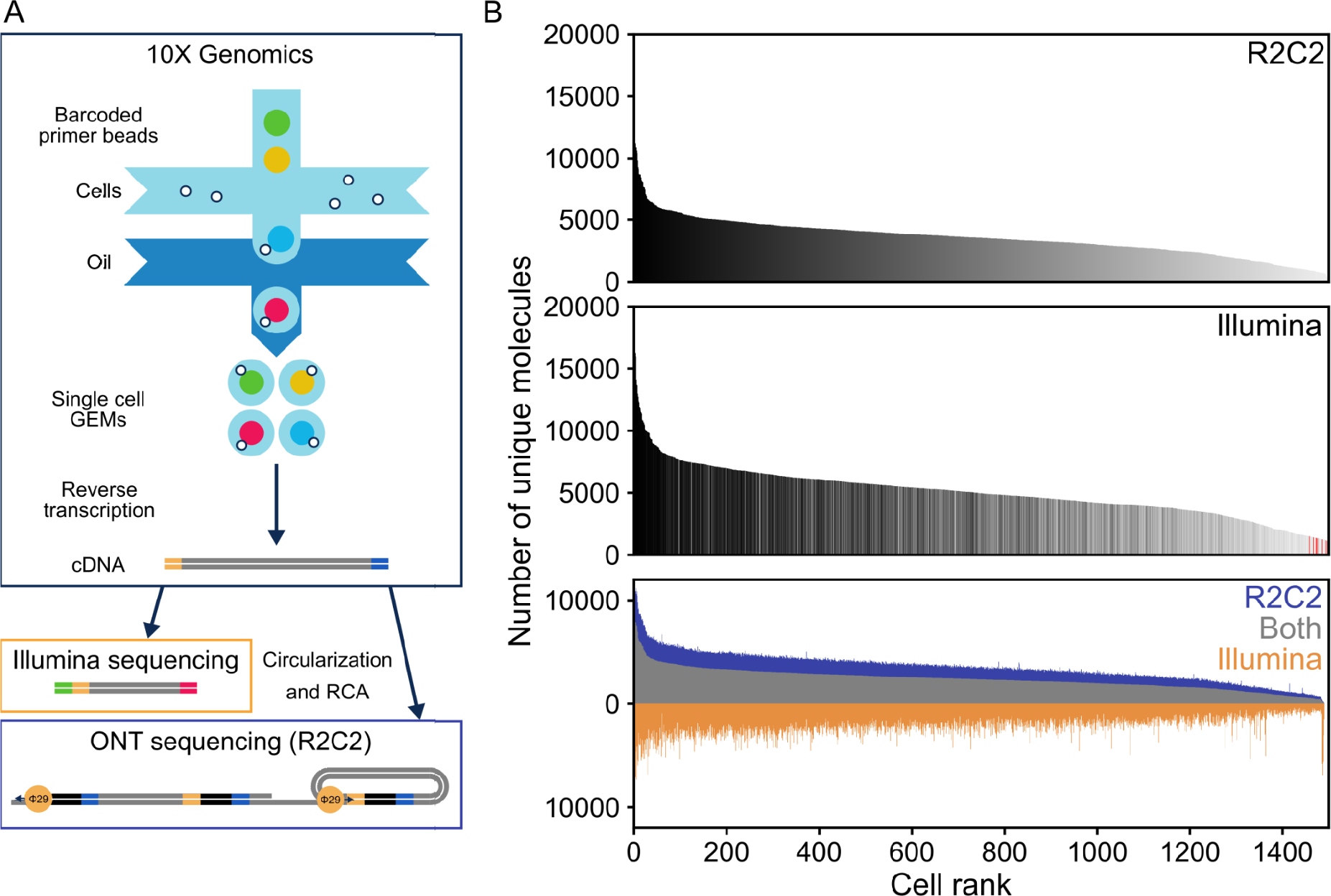
Data Generation and Characteristics. A) Thousands of peripheral blood mononuclear cells (PBMCs) were processed using the 10X Genomics Chromium Single Cell 3’ Gene Expression Solution. The resulting full-length cDNA was either fragmented for Illumina sequencing or processed using the R2C2 workflow. B) After read processing and demultiplexing, the unique molecular identifiers (UMIs) associated with each cellular index (cell) in R2C2 (top) and Illumina (center) datasets are shown as histograms. Cells are ranked by the number of UMIs and colored based on their rank in the R2C2 dataset. Red lines indicate cellular identifiers found in Illumina but not R2C2 data. At the bottom, the UMIs shared between cellular identifiers in Illumina and R2C2 datasets or unique to each dataset are shown as stacked histograms. Cells are ranked by the number of shared UMIs. Data for replicate 1 are shown.

**Fig. 2:**
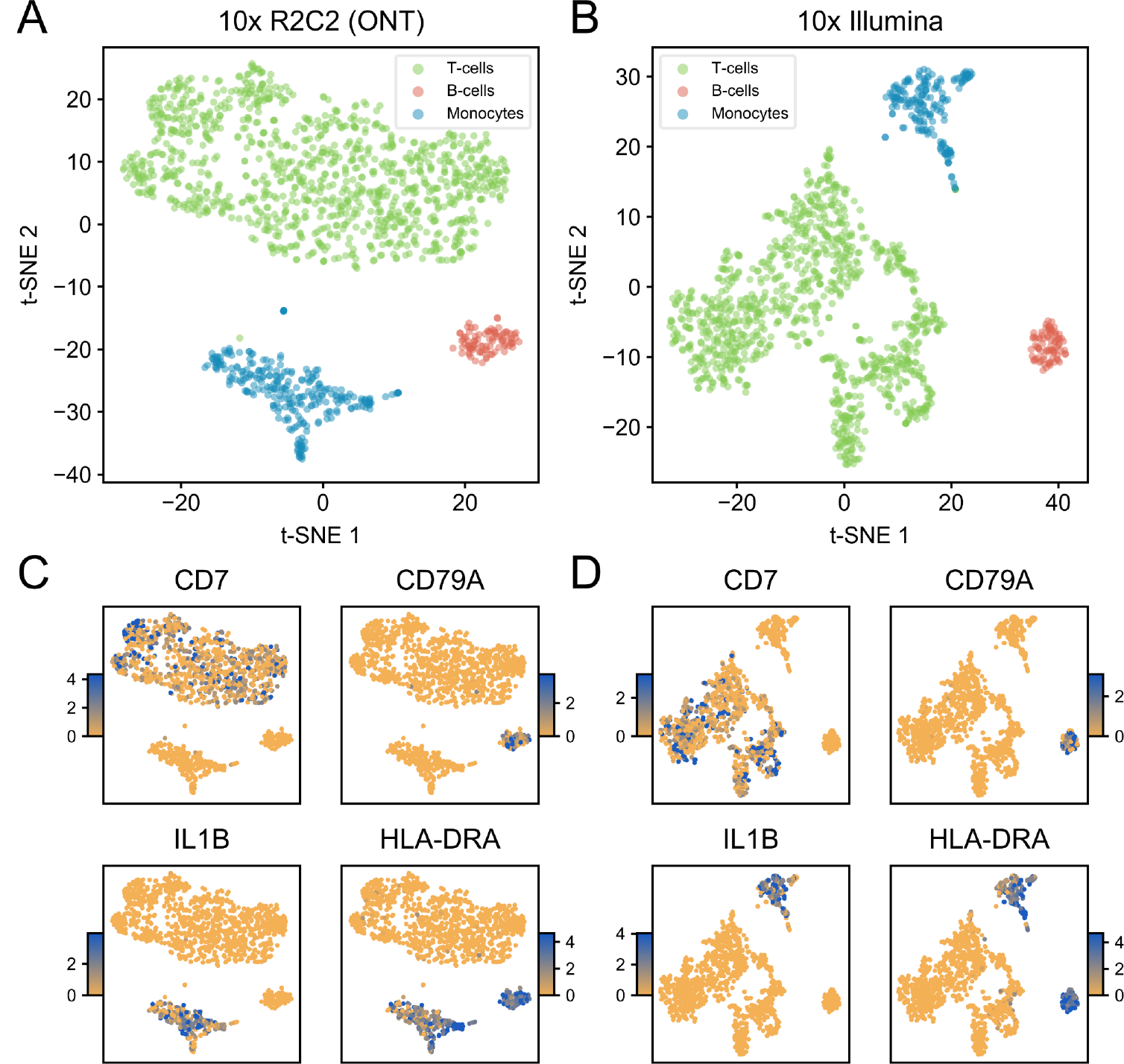
R2C2 and Illumina datasets independently cluster into B cells, T cells, and Monocytes. Gene expression profiles were determined independently for each cell in R2C2 and Illumina datasets. The Seurat package was then used to cluster cells based on the gene expression profiles. The cells in R2C2 (A) and Illumina (B) datasets both clustered into 3 groups which, based on marker gene expression (C and D) could be identified as B cells, T cells, and Monocytes. The color gradient (C and D) encodes ln(fold change), where the fold change is comparing that cluster’s expression to the rest of the data. Data for replicate 1 are shown.

### Illumina data covers 10X UMIs comprehensively

For sequencing on the Illumina NextSeq, we fragmented the full-length cDNA according to the standard 10X protocol. We demultiplexed and merged the resulting reads based on cellular barcodes and unique molecular identifiers (10X-UMIs) associated with every amplified transcript molecule during reverse transcription (see Methods). By only keeping transcript molecules with a raw read coverage of >3, we condensed 202,469,707 raw read pairs to 15,264,862 reads originating from the 3’ ends of unique transcript molecules across both replicates (∼5000 molecules per cell).

### R2C2 data identifies the same cellular and molecular identifiers as Illumina data

For sequencing on the ONT MinION and PromethION sequencers, we processed 10ng of full-length cDNA using the previously published R2C2 workflow (see Methods). The resulting R2C2 libraries were then sequenced using standard ONT LSK-109 ligation based sequencing kits. We processed the resulting ONT raw reads into R2C2 consensus reads using the C3POa pipeline (Table 1 and S1). We then merged reads in two sequential steps if they contained matching unique molecular identifiers (UMIs) in either the dsDNA splint used to circularize cDNA molecules (Splint-UMI) or the 10X oligo(dT) primer used to prime reverse transcription of poly(A) RNA molecules (10X-UMI).

**Table 1:** Read numbers throughout processing.

|  | Basecalled reads | Post-processed R2C2 consensus reads | Splint-UMI merged R2C2 consensus reads | Splint/10X-UMI merged R2C2 consensus reads | Demultiplexed R2C2 reads |
| --- | --- | --- | --- | --- | --- |
| **Replicate 1** | 29,529,179 | 11,564,494 (39.2%) | 11,368,091 (98.3%) | 7,853,440 (69.1%) | 6,385,901 (81.3%) |
| **Replicate 2** | 26,526,607 | 10,661,139 (40.2%) | 10,276,420 (96.4%) | 6,968,632 (67.8%) | 5,652,620 (81.1%) |

First, we merged 3.3% and 6.5% of the R2C2 consensus reads in replicate 1 and replicate 2 respectively because their Splint-UMI identified them as originating from the circularization of the same cDNA molecule. Second, we merged 46.3% and 46.1% of these Splint-UMI merged R2C2 consensus reads in replicate 1 and replicate 2, respectively, because their 10X-UMI identified them as originating from the same RNA molecule. Across both replicates this sequential merging process resulted in 14,822,072 Splint/10X-UMI merged R2C2 consensus reads (Table S2) with a median sequence accuracy of 98.0%.

Next, we demultiplexed these ∼14.8 million Splint/10X-UMI merged R2C2 consensus reads based on the 10X cellular barcodes they contained. In this way, 81% of these reads could be successfully assigned to an individual cell, which compares favorably to the ∼6% Illumina-independent and ∼67% Illumina-guided assignment rates determined for standard ONT reads in previous studies [24,29].

Moreover, 2974 (99.1%) of the 3000 cellular identifiers we determined independently from the R2C2 dataset also appeared in the Illumina dataset.

Because we merged reads in Illumina and R2C2 datasets based on the 10X-UMI, each read in either dataset should originate from a unique RNA molecule. Consequently, the number of reads assigned to each cell was also highly similar between the datasets (Fig 1B). Also, for each cell, 67% of the R2C2 reads contained a 10X-UMI that was also present in an Illumina read assigned to the same cell. Interestingly, the accuracy of R2C2 reads containing 10X-UMIs present in an Illumina read was significantly higher than the accuracy of R2C2 reads containing 10X-UMIs not present in an Illumina read (98.4% vs. 97.1%; p=0.0 Monte-Carlo Permutation test). This indicates that read accuracy plays an important role in accurately identifying UMI sequences. However, although their RNA molecule of origin cannot be unambiguously identified, R2C2 reads containing UMIs with sequencing errors are still highly valuable for downstream analysis.

### Clustering single cells into cell types based on gene expression

We next investigated whether these R2C2 reads could be used to determine gene-expression accurately enough to cluster single cells into cell types – an analysis step that is currently routinely performed using short-read based gene expression. To this end, we used minimap2 to align R2C2 reads to the human genome (hg38) and used featureCounts to determine gene expression levels in each cell [30,31]. For comparison, Illumina reads generated from the same cDNA were aligned using STAR and also processed using featureCounts [32]. Median Pearson-r values for R2C2 and Illumina-based gene expression for the same cell showed high correlation at 0.74 (Fig. S1).

We then clustered R2C2 and Illumina datasets independently using the Seurat analysis package [33]. R2C2 and Illumina datasets both grouped into three cell type clusters. Based on marker gene expression, the major cell types could be identified as B cells (CD79A) [34], T cells (CD7) [35], and Monocytes (IL1B) [36] – the expected composition of a PBMC sample (Fig. 3, S2). Importantly 99.4% of cells that were clustered in both datasets associated with the same cell type in the two datasets.

**Fig. 3:**
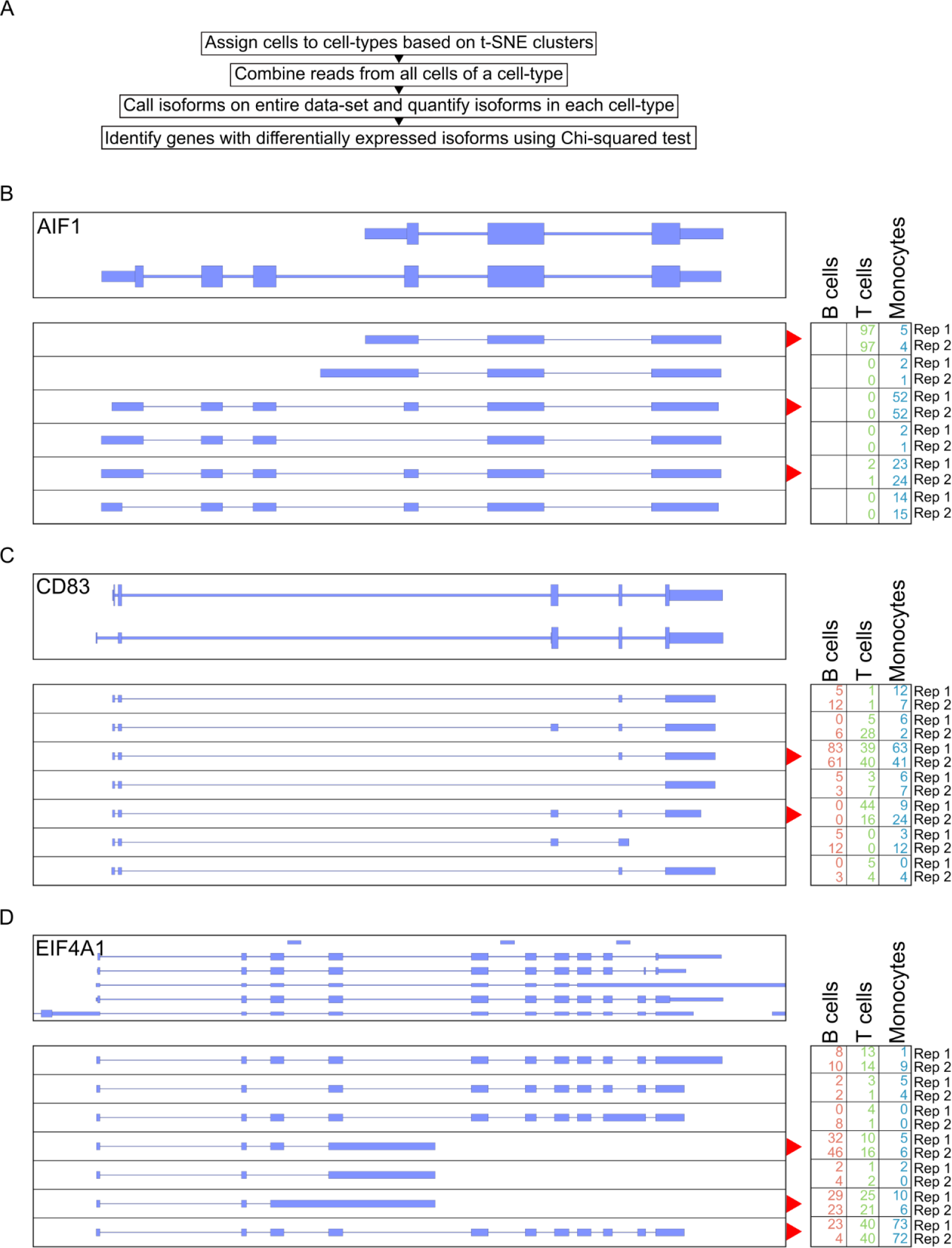
Identifying differentially expressed isoforms between cell types using clustered single cell data. **A)** Workflow of differentially expressed isoform identification. R2C2 reads are separated by cell type, then used to identify and quantify isoforms. Genes with differential isoform usage between cell types are then identified using Chi-squared tests. B-D) Genome Browser shots of three genes with differential isoform expression. Gene annotation is shown on top. Isoforms as determined by Mandalorion on the entire dataset are shown below (“top strand”=blue, “bottom strand”=yellow). Relative quantification (%) of each isoform in each cell type and replicate is shown on the right. Isoforms with the most variable changes in abundance are indicated with a red arrow.

This showed that R2C2 reads show performance comparable to Illumina data for determining gene expression and clustering cell types in massively multiplexed single-cell experiments.

### Generating cell type specific isoform-level transcriptomes

Having successfully sorted cells into cell types, we set out to generate high quality transcriptomes for these cell types. This is possible because, as shown in previous studies analyzing 10X cDNA with long reads [24,26], R2C2 reads appeared to cover entire transcripts (Fig. S3)

First, as previously established [26], we pooled all reads associated with the cells of each cell type to create a synthetic bulk sample. We then identified transcript isoforms for each synthetic bulk cell type using Mandalorion [19–21,27]. The majority (50-60%) of isoforms generated by Mandalorion for the individual cell types were classified by SQANTI [37] as either ‘full-splice-match’ or ‘novel-in-catalog’, which represent likely full-length isoforms. This number increased to >80% if only multi-exon isoforms were considered. In aggregate, the cell type specific isoforms we generated represent full-length B cell, Monocyte, and T cell transcriptomes, with each transcriptome’s depths dependent on the number of cells and reads associated with each cell type (Table 2). With ∼9 million R2C2 reads and 14,925 multi-exon isoforms, the T cell transcriptome is the most complete and likely most useful of the three cell types.

**Table 2:** Cell type specific full-length transcriptome characteristics.

| cell type | Number of cells | Number of reads | Number of genes with multi-exon isoforms | Number of multi-exon isoforms |
| --- | --- | --- | --- | --- |
| B cells | 179 | 625,334 | 1,481 (plus 55 novel genes) | 2006 |
| T cells | 2,199 | 9,108,828 | 6,934 (plus 448 novel genes) | 14,925 |
| Monocytes | 464 | 2,042,162 | 2,882 (plus 77 novel genes) | 4530 |

### Differential isoform usage between cell types

In addition to determining which isoforms are expressed, we can also quantify the expression of these isoforms and investigate whether they are differentially expressed between the three cell types. To perform this differential isoform expression analysis, we first wanted to capture all the isoforms expressed in the entire dataset. To this end, we composed an additional synthetic bulk sample using the R2C2 reads from all cells in the dataset. We then used Mandalorion to identify all isoforms present in this synthetic bulk sample and quantified the expression of each isoform in B cells, T cells, and Macrophages. Next, quantified isoforms were grouped by the genes they were associated with and genes with significant isoform usage between cell types were determined using a Chi-square contingency table test. After filtering for genes expressed in at least two cell types and multiple testing correction, we identified 74 genes with differential isoform usage (p-value<0.01)(Table S3). The features that distinguished differentially expressed isoforms included alternative TSSs (AIF1, Fig. 3B), cassette exons (CD83, Fig. 3C), or poly(A) sites (EIF4A1, Fig. 3D).

### Isoform diversity is highly variably between genes

Next we investigated whether single-cell derived transcriptome information can enrich our understanding of isoform diversity. While pooling all reads associated with a cell type can serve as a basis for defining transcriptome annotations, this approach loses information on which isoforms are expressed by which individual cell and due to coverage cut-offs likely presents a conservative estimate of the true isoform diversity present in a cell type.

In the 3000 cell dataset we present here, we have sufficient coverage to generate isoforms for each cell independently. Using Mandalorion, we generated a median of 127 multi-exon isoforms per cell, with the majority being classified as either ‘full-splice-match’ (77%) or ‘novel-in-catalog’ (11%).

We then analyzed isoform diversity across ∼3000 the cells in the dataset. To this end, we merged identical isoforms expressed by different cells. We then determined how many cells expressed isoforms for any given gene.

Interestingly, isoform diversity varied greatly between genes (Fig. 4A). On one end of the spectrum, genes encoding ribosomal proteins in particular are expressed in the majority of cells, yet we identify few unique isoforms for these genes. For example, 1299 cells expressed a total of 1299 isoforms (as determined by Mandalorion) of the ribosomal protein gene RPL35. After merging all identical isoforms, only 8 unique isoforms remained and only one of those was expressed by more than one cell. On the other end of the spectrum, genes like LMNA are also expressed by a majority of cells but feature many unique isoforms. In fact, 930 cells expressed a total of 969 unique LMNA isoforms. After merging all identical isoforms, only 305 unique isoforms remained and 86 of those were expressed by more than one cell.

**Fig. 4:**
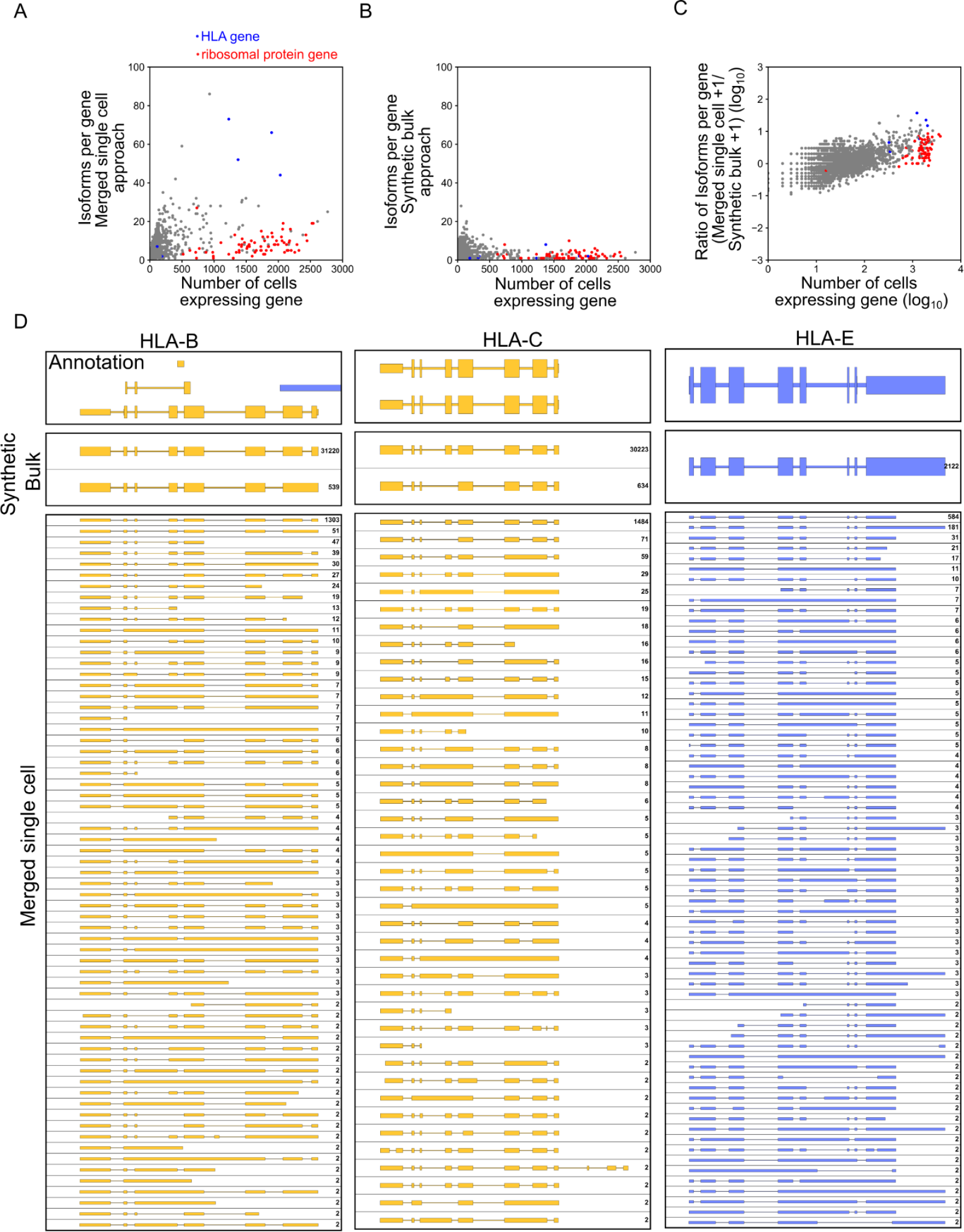
Genes show a wide range of isoform diversity. We generated an isoform level transcriptome for each cell in our dataset and then analyzed the isoform diversity for different genes by merging these isoforms. A) The correlation of the number of cells expressing an isoform for a gene and how many unique isoforms we identified for that gene using the ‘merged single cell’ approach is shown as a scatter plot. B) The correlation of the number of cells expressing an isoform for a gene and how many unique isoforms we identified for that gene using the synthetic bulk approach is shown as a scatterplot. C) The correlation of the number of cells expressing an isoform for a gene the ratio of the number of isoforms identified for that gene with the ‘merged single cell’ and ‘synthetic bulk’ approaches. Both number of cells and isoform ratio are shown as log_10_. ABC) Genes encoding ribosomal proteins and HLA proteins are shown in red and blue respectively. D) Genome Browser shots HLA genes are shown. Genome annotation is shown on top, isoforms determined by the synthetic bulk approach in the middle, and isoforms determined by the merged single cell approach at the bottom. (“top strand”=blue, “bottom strand”=yellow). Number of reads (synthetic bulk) or cells (merged single cells) associated with an isoform are shown on the right.

Unique isoforms expressed by more than one cell as determined by this *‘merged single cell’* approach could therefore be used to enrich isoform annotations based on bulk or synthetic bulk data. For example, combining all R2C2 reads collected for all the cells in this study and identifying isoforms based on this synthetic bulk yielded one isoform for RPL35 but also only 3 isoforms for LMNA, likely due to minimum relatives abundance requirements of 1% at a locus set as default in Mandalorion. In fact, most genes expressed by many cells had a low number of isoforms identified by the ‘*synthetic bulk’* approach (Fig. 4B).

By systematically comparing the number of isoforms determined by *‘merged single cell’* and ‘*synthetic bulk*’ approaches we showed that the more cells expressed isoforms for a gene, the more likely the *‘merged single cell’* approach was to identify additional isoforms. This analysis highlighted the behavior of HLA class I genes, in particular HLA-B, HLA-C, and HLA-E (Fig. 4C), which all showed >40 isoforms with the *‘merged single cell’* approach but only one or two in the *‘synthetic bulk’* approach (Fig. 4A,B,D).

### Extracting paired adaptive immune receptor sequences from B and T cells

In addition to the analysis of regular transcript isoforms, we investigated whether our datasets enable the identification and pairing of adaptive immune receptor (AIR) transcripts. AIR transcripts encode for antibodies and T cell receptors which pose unique challenges for sequencing applications. Each antibody (IG) or T cell receptor (TR) is encoded by two AIR transcripts each of which is transcribed from a gene whose V (, D,) and J segments are uniquely rearranged in each individual B or T cell.

Our standard Mandalorion transcript isoform identification workflow does not capture these AIR transcripts reliably because it relies on read alignments which fail for the highly repetitive and rearranged IG heavy (IGH), IG light (IG kappa (IGK) and lambda (IGL)), TCR alpha (TRA), and beta (TRB) loci. To capture AIR transcripts reliably, we first identified R2C2 reads which aligned to the constant region exons in the IG and TR loci. We then determined which of these reads contained a high quality V segment using IgBlast [38]. Finally, we used these filtered reads to determine consensus sequences for each locus and cell (Fig. 5A).

**Fig. 5:**
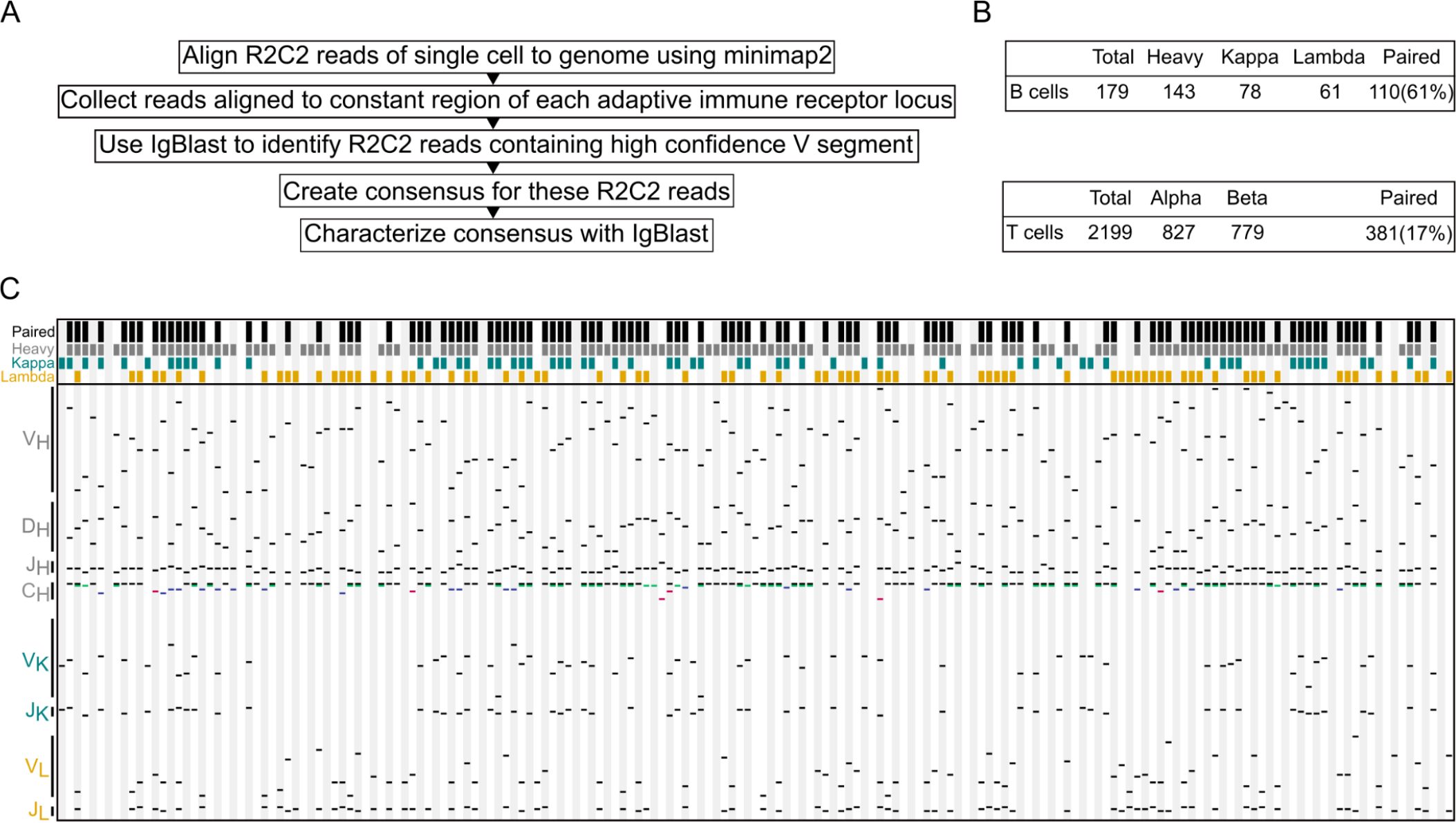
IG and TCR transcripts can be identified and paired in 10X R2C2 data. A) The workflow to identify antibody (IG) and T cell receptor (TCR) transcripts for each individual cell. B) Numbers of cells for which IG or TCR transcripts could be identified and paired. C) Schematic of IG identification, composition, and pairing. Each column represents a single B cell. Colored blocks on top of each column indicate whether a cell contains paired IG transcripts (black), whether an IGH (Heavy: grey), IGK (Kappa: teal), or IGL (Lambda: orange) transcripts was detected. Below the diversity of the detected sequences is shown. Black lines indicate which gene segments were used when an IG sequence was recombined from the germline genome. In C_H_, it is also shown which isotype(s) we detected (IGHM: black, IGHD: green, IGHA1 or 2: red, IGHG1-4: blue) for each cell.

For many B cells we determined multiple sequences for different isotypes (IGHM, IGHD, IGHG(1, 2, 3,and 4), and IGHA(1 and 2)) (Table S4) and isoforms (membrane bound and secreted). In the vast majority of cases (103/108) (Fig. 5B), transcripts contained the same V segment, indicating that they represent alternative splicing products of the same rearrangement. We succeeded in determining paired IG sequences for 110 B cells and 381 T cells which represent 61% and 17% of all B and T cells analyzed in this study, respectively (Fig. 5C). Importantly, as would be expected for a random sample of B cells, the V (, D,) and J segment usage composition of the paired transcripts of these cells was highly diverse (Fig. 5C)

## Discussion

Here, we present a method to analyze highly-multiplexed full-length single-cell transcriptomes that does not require short-read sequencing. We processed 10ng of cDNA generated as an intermediate product of the 10X Genomics Chromium Single Cell 3’ Gene Expression Solution into R2C2 sequencing libraries. We sequenced these libraries and demultiplexed the resulting data to produce over 12 million unique transcript molecules generated from ∼3000 PBMCs. We used these single cell data to determine monocyte, T cell, and B cell clusters, generate isoform-level transcriptomes for these cell types, investigate single-cell isoform diversity, and pair adaptive immune receptor transcripts.

The ability to analyze the full-length transcriptomes of single cells without the need for Illumina short-read data has the potential to simplify experimental workflows. The ability to perform this analysis on low cost ONT sequencers will make it more accessible. This is made possible through the use of the R2C2 sample preparation method which can increase the base accuracy of ONT MinION sequencers to ∼99%. In this study, the R2C2 base accuracy was closer to 98% due to shorter raw reads. We aimed for shorter raw reads to increase R2C2 read numbers and, to this end, reduced the stringency of our size-selection prior to sequencing (Table S1).

Outside of R2C2, raw nanopore reads are becoming more accurate and are used to analyze 10X cDNA with the help of Illumina data or by themselves using modified 10X protocols with longer indexes. Further, single cell studies using the PacBio Sequel II, while limited in overall throughput and hampered by per-read cost of the sequencer, benefit from the very high accuracy of the reads which simplifies computational analysis. Going forward, the trade-off between throughput, cost, and accuracy of ONT MinION and PromethION as well as PacBio Sequel II sequencers will have to be considered closely and the best compromise may well vary between studies.

At current throughput and accuracy, the combination of ONT sequencers and the R2C2 method allows the analysis of thousands of cells. An increase in read output will make it possible to either analyze more cells or sequence all transcripts reverse transcribed by the 10X Genomics workflow. In this current study, with about 3000 R2C2 reads per cell, we captured about 67% of the molecules present in an exhaustively sequenced Illumina dataset of the same cDNA. This was sufficient to cluster cell types and generate single-cell transcriptomes. An increase in accuracy would make future demultiplexing and UMI merging steps more efficient. While our demultiplexing strategy can handle sequencing errors (see Methods), at 98% accuracy it still only manages to demultiplex ∼81% of R2C2 reads, which is better than previously published approaches, but not ideal [24,29]. Increasing accuracy could increase this number significantly. Paired with higher throughput, future experiments could only retain UMIs which were observed more than once, similar to how we analyze Illumina data (see Methods).

Beyond establishing this method, we generated high-quality transcriptomes for Monocyte, B cell, and T cell populations. Because the majority of PBMCs are T cells, the T cell transcriptome is the most comprehensive of those three and should serve as a resource for understanding the biology of these adaptive immune cells.

We then used a framework developed for a previous study [28] to show that these cell types show differential isoform expression. The ability to identify differentially expressed isoforms expands the quality of information that can be extracted from single-cell experiments and opens the door to a much more nuanced understanding of gene regulation.

Beyond investigating isoform expression on the cell type level, we investigated the extent of isoform diversity on the single-cell level. While some genes showed low isoform diversity, i.e. most cells express the same isoform, some genes showed high diversity, i.e. many cells express unique isoforms. This wide range of isoform diversity will pose a formidable challenge for single-cell level differential isoform expression analysis going forward. Future studies into how this wide range of isoform diversity is maintained and used by cells are bound to generate fascinating insights into transcript processing and cellular function.

In the meantime, using isoforms identified independently for single cells can already inform isoform identification. While different isoform identification tools like TALON [39], FLAIR [40], or StringTie2 [41], and Mandalorion use different strategies when identifying and filtering isoforms, they all rely on some form of read coverage cut-off to differentiate real isoforms from the noise produced by any sequencing method. However, PCR or sequencing artifacts generated within a single cell can overcome these cut-offs and result in the false-positive identification of isoforms. The information of how many single cells express an isoform could therefore aid in the identification of real or biologically meaningful isoforms as each single cell can be seen as an independent biological replicate.

Finally, taking advantage of the single-cell nature of this dataset, we performed analysis on the most complex part of T cell and B cell transcriptomes, namely adaptive immune receptor transcripts. By sequencing and pairing adaptive immune receptor transcripts expressed by single T and B cells, we showcased the power of long reads for resolving even the most challenging transcript isoforms – without the need for specialized protocols. This will be of particular use when analyzing complex samples that contain, but aren’t limited to, immune cells like solid or liquid tumors.

## Methods

### Single cell cDNA library preparation

Full-length cDNA pools and Illumina libraries were prepared by 10X Genomics. PBMCs were sourced from Stemcell Technologies and prepared for sequencing using the 10X Genomics Chromium Single Cell 3’ Gene Expression Solution. Preparation of the cDNA was done according to manufacturer’s instructions with the exception of the extension time for the final PCR reaction which was standard 1 minute for replicate 1 but increased to 4 minutes for replicate 2.

### Illumina sequencing and read processing

Illumina libraries were sequenced on the Illumina NextSeq with Read1 = 26bp and Read2 = 134bp.

Overall a NextSeq flowcell generated 107,911,006 reads for replicate 1 and 75,753,410 reads for replicate 2. Reads were then demultiplexed and collapsed by determining the 1500 most frequent cellular barcodes, perfectly matching cell barcodes to the most frequent, and then filtering for unique cell barcode/10X UMI combinations.

Reads for each cell were then aligned to the human genome (hg38) using STAR (*--runThreadN 30 –genomeDir /path/to/STAR/index/ --outSAMtype BAM SortedByCoordinate --readFilesIn /path/to/reads –outFileNamePrefix /path/to/alignment/dir*).

### Nanopore sequencing and read processing

Full-length cDNA pools were prepared as described previously. In short, 10ng of cDNA is circularized using a DNA splint compatible with 10X cDNA and the NEBuilder HIFI DNA Assembly Master Mix (NEB). The DNA splint was generated by primer extension of the following oligos:

~~~
>10X_UMI_Splint_Forward (Matches 10X PCR primer)
AGATCGGAAGAGCGTCGTGTAG
TGAGGCTGATGAGTTCCATANNNNNTATATNNNNNATCACTACTTAGTTTTTTGATAGCTTCAAGCCAGAGTTGTCTTTTTCTCTTTGCTGGCAGTAA
AAG
>10X_UMI_Splint_Reverse (Matches ISPCR Primer)
CTCTGCGTTGATACCACTGCTT
AAAGGGATATTTTCGATCGCNNNNNATATANNNNNTTAGTGCATTTGATCCTTTTACTCCTCCTAAAGAACAACCTGACCCAGCAAAAGGTACACAAT
ACTTTTACTGCCAGCAAAGAG
~~~

Non-circularized DNA is digested using Exonucleases I, III, and Lambda. Circularized DNA is amplified using rolling circle amplification using Phi29 (NEB). The resulting HMW DNA is debranched using T7 Endonuclease (NEB) and purified and size-selected using SPRI beads. This DNA containing concatemers of the originally circularized cDNA is then sequenced using the LSK-109 kit on either ONT MinION or PromethION sequencers (Table S1). The resulting raw reads were processed into consensus reads using the C3POa pipeline (v2.2.2). All consensus reads were then assigned a cell of origin. In a first step, we determined the most common ∼1500 cellular identifiers in our sample using a simple counting strategy. Then, we assigned reads to the most similar cellular identifiers if they fit the following criteria:

1. *L1* < *3* and
2. *L1* < *L2 - 1*

*where*

*L1 is the Levenshtein distance between the read’s cellular identifier and the most similar known cellular identifier*

*and*

*L2 is the Levenshtein distance between the read’s cellular identifier and the second most similar known cellular identifier*.

These consensus reads were demultiplexed based on their cell assignment, they were merged if the contained the similar UMIs in their splint back-bones using the ExtractUMIs and MergeUMIs utilities (https://github.com/rvolden/10xR2C2). The resulting reads were then merged again if they contained the similar 10X UMIs in their adapters using the ExtractUMIs and MergeUMIs utilities (https://github.com/rvolden/10xR2C2).

The resulting Splint/10X-UMI merged R2C2 consensus reads were then demultiplexed based on their initial cell assignments. If a Splint/10X-UMI merged R2C2 consensus read was generated by merging reads with different cell assignments it was discarded. Reads for each cell were then aligned to the human genome (hg38) using minimap2 [30] (*-ax splice --secondary=no -G 400k*).

### Cell type clustering

Both Illumina and R2C2 data were analyzed in the same way independently. First gene expression tables were generated using featureCounts [31]. Then these tables were parsed for input into the Seurat R package (v3) [33]. Seurat generated cell type clusters using the following main settings (*min*.*cells=3, min*.*features=200, percent*.*mt<5, 2500>nFeature_RNA>200, nfeatures=2000, dims=1:10, resolution=0*.*08 (0*.*08 used for nanopore, 0*.*03 for Illumina), log normalization, and vst selection*).

For each cell, cell type information was extracted based on location for downstream analysis.

### Isoform analysis

We generated high confidence isoforms using the latest version of the Mandalorion pipeline (Episode III.5, https://github.com/rvolden/Mandalorion).

#### Cell type transcriptomes

All reads and subreads assigned to cells of a cell type were pooled. Mandalorion was run on these files with the following settings:

~~~
-c /path/to/config_file
-m /path/to/NUC.4.4.mat
-I 300
-g /path/to/gencode.v37.annotation.gtf
-G /path/to/hg38.fa
-a /path/to/10x_Adapters.fasta
-f /path/to/Pooled_reads.fa
-b /path/to/Pooled_subreads.fa
-p /path/to/output_folder
-e ATGGG,AAAAA
~~~

with `10x_Adapters.fasta` containing the following sequences:

~~~
>3Prime_adapter
CTACACGACGCTCTTCCGATCT
>5Prime_adapter
AAGCAGTGGTATCAACGCAGA
~~~

#### Single-cell transcriptomes

Mandalorion was run on the reads, read alignments, and subreads of each individual cell. Mandalorion was run with the following settings:

~~~
-c /path/to/config_file
-I 300
-g /path/to/gencode.v37.annotation.gtf
-G /path/to/hg38.fa
-a /path/to/10x_Adapters.fasta
-f /path/to/SingleCell_reads.fa
-b path/to/SingleCell_subreads.fa
-p path/to/output_folder
-e ATGGG,AAAAA
-R 2
~~~

Note that we reduced the minimum number of reads required to identify an isoform to 2.

The resulting isoform psl files were converted to gtf files and classified using the sqanti_qc.py program and the following settings:

~~~
-g
-n
-t 24
-o output_prefix
-d /path/to/output_folder
path/to/gtf_file /path/to/gencode.v37.annotation.gtf /path/to/hg38.fa
~~~

#### Isoform diversity analysis

Similar isoforms were merged using the `merge_psls.py` utility which accepts a list of isoform fasta and psl files and merges isoforms if they:

1. Use all the same splice sites This step is base-accurate but will treat splice site a single base pair apart as equivalent if one site is much less abundant than the other
2. Use the similar start and end sites

This step will consider sites similar if they are at most 10nt apart. Because isoforms are iteratively grouped at this step, individual isoforms in a merged group might have sites that are further than 10nt apart but are connected by a third isoform between them.

### Adaptive Immune receptor analysis

For each cell, reads aligning to the T cell or B cell receptor loci were extracted from sam files using samtools view [42] and the below genomic coordinates.

~~~
IGH:-chr14:-105,533,853---106,965,578
IGK:-chr2:-89,132,108---90,540,014
IGL:-chr22:-22,380,156---23,265,691
TRA:-chr14:-22,178,907---23,021,667
TRB:-chr7:-141,997,301---142,511,567
~~~

Reads were then analyzed for each cell and locus (and for IGH, each isotype/isoform) separately by filtering reads for a high-quality match to a V segment retrieved from IMGT [43] using IgBlast [38] and the following settings:

~~~
-germline_db_V /path/to/V_segments
-germline_db_J /path/to/J_segments
-germline_db_D /path/to/D_segments
-organism human
-query /path/to/reads.fasta
*[-ig_seqtype TCR] - only for T cell receptors*
-auxiliary_data optional_file/human_gl.aux
-show_translation
-outfmt 19
~~~

Filtered reads for each cell were then used to generate consensus reads for each locus. Those consensus reads were then assigned V, (D,) and J segments using IgBlast and the same settings as above. All scripts used for this analysis and a wrapper script automating this analysis are available at https://github.com/christopher-vollmers/AIRR-single-cell.

### Data Access

We uploaded all data generated for this study to the SRA where it is available under BioProject accession PRJNA599962.

B cell, T cell, and Monocyte transcriptomes are available at https://users.soe.ucsc.edu/~vollmers/10XR2C2/.

### Code Access

We have made the code required to demultiplex R2C2 reads and format gene expression matrices for Seurat available on GitHub (https://github.com/rvolden/10xR2C2). Code for AIRR analysis is also available on GitHub (https://github.com/christopher-vollmers/AIRR-single-cell).

## Acknowledgements

We thank 10X Genomics for generating full-length cDNA and Illumina sequencing libraries from human PBMCs. We acknowledge funding by the National Human Genome Research Institute/National Institute of Health Training Grant 1T32HG008345-01 (to R.V.), the Hellman Foundation, Santa Cruz Cancer Benefit Group, and National Institute of General Medical Sciences/National Institute of Health Grant R35GM133569 (to C.V.)

## Supplementary Information to

**Supplementary Figure S1:**
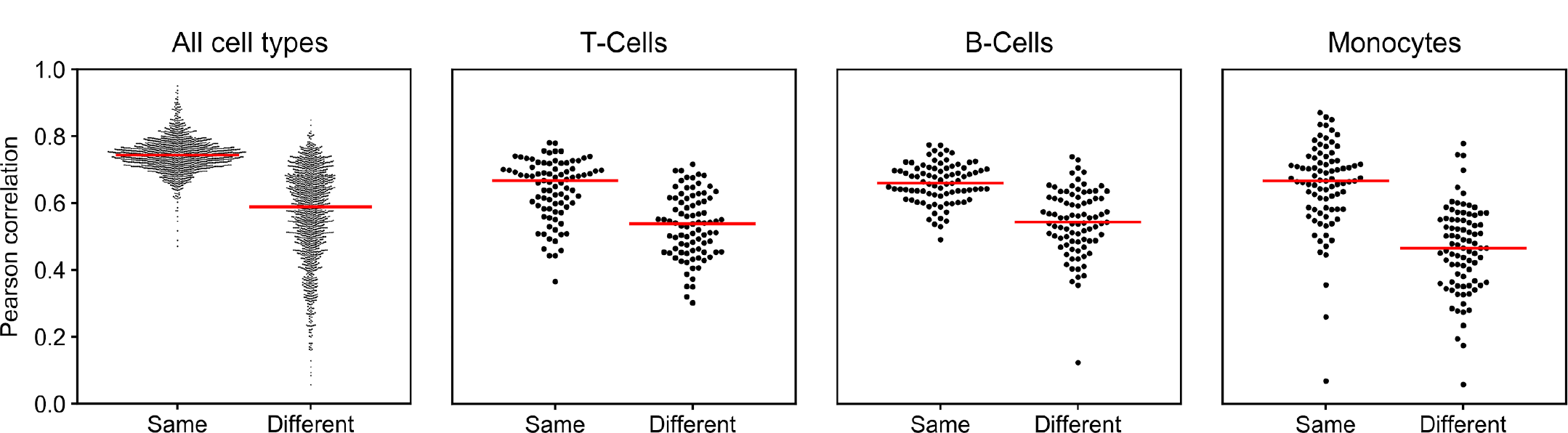
Swarm plots of gene expression correlation between R2C2 and Illumina. The median Pearson correlation for each swarm is shown in red. From left to right: (All cell types, same) cells were matched based on their cellular barcode from R2C2 and Illumina. (All cell types, different) R2C2 cells were correlated to a random cell in the Illumina data. The next three swarms were subsampled to 85 points because there are 89 B-Cells. (T-Cells, same) Random T-Cells were correlated between R2C2 and Illumina data. (T-Cells, different) Random R2C2 T-Cells were correlated with random Illumina non-T-Cells. (B-Cells, same) Random B-Cells were correlated between R2C2 and Illumina. (B-Cells, different) Random R2C2 B-Cells were correlated with random Illumina non-B-Cells. (Monocytes, same) Random Monocytes were correlated between R2C2 and Illumina. (Monocytes, different) Random R2C2 Monocytes were correlated with random Illumin non-Monocytes.

**Supplementary Figure S2:**
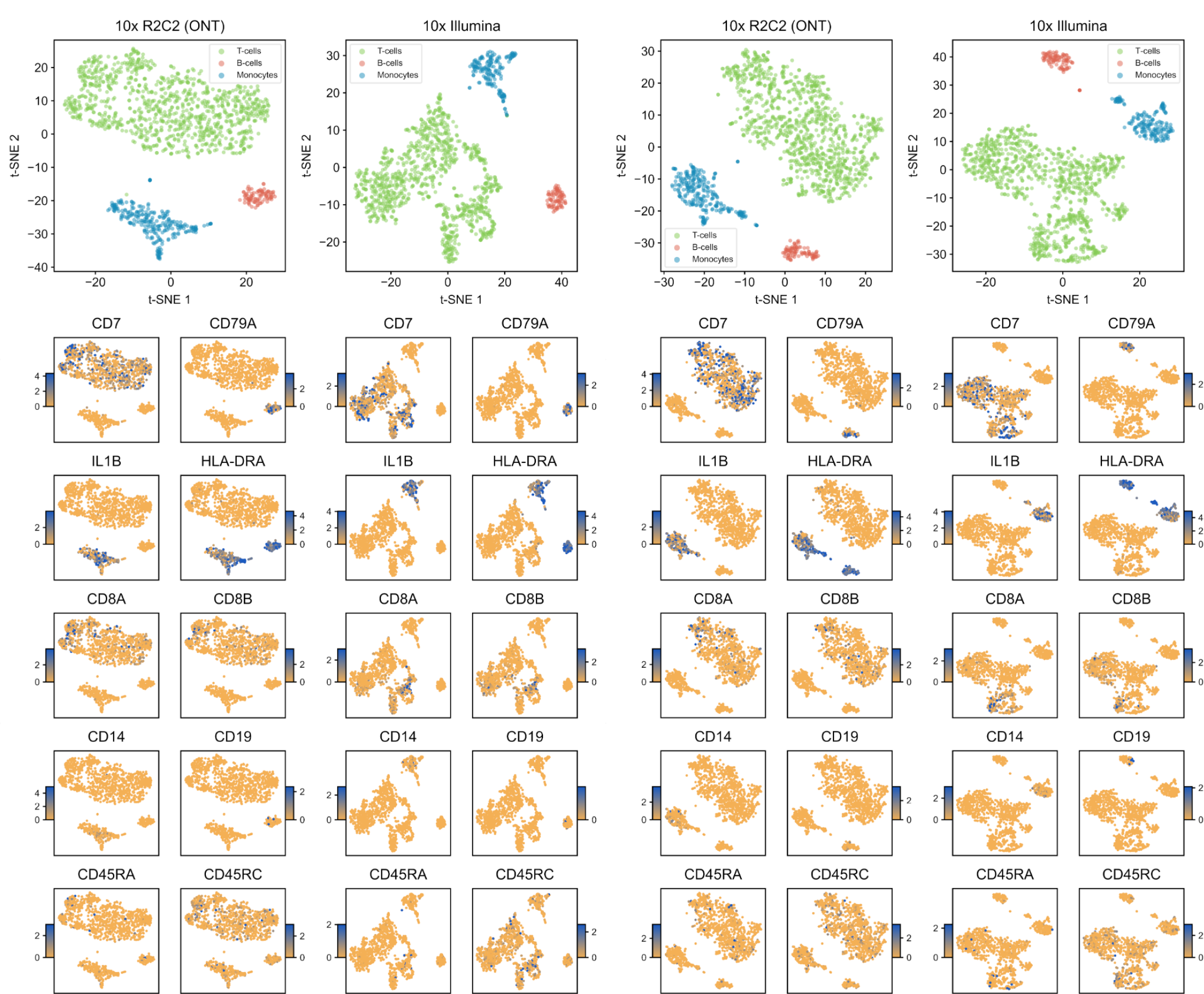
t-SNE plots with additional marker genes for replicates 1 and 2. As for Figure 2 plots are based on gene expression data as calculated by featureCounts and Seurat. Plots for replicate 1 and replicate 2 are shown on the left and right respectively. Top left: replicate 1 cell type clusters for R2C2 and Illumina. Bottom left: replicate 1 expression heat maps for various marker genes where the two columns on the left are for R2C2 and the right two are Illumina. Top right: replicate 2 cell type clusters for R2C2 and Illumina. Bottom right: replicate 2 expression heat maps for various marker genes where the two columns on the left are for R2C2 and the right two are Illumina. Additional marker genes taken from [14]. The color gradient encodes ln(fold change), where the fold change is comparing that cluster’s expression to the rest of the data.

**Supplementary Figure S3:**
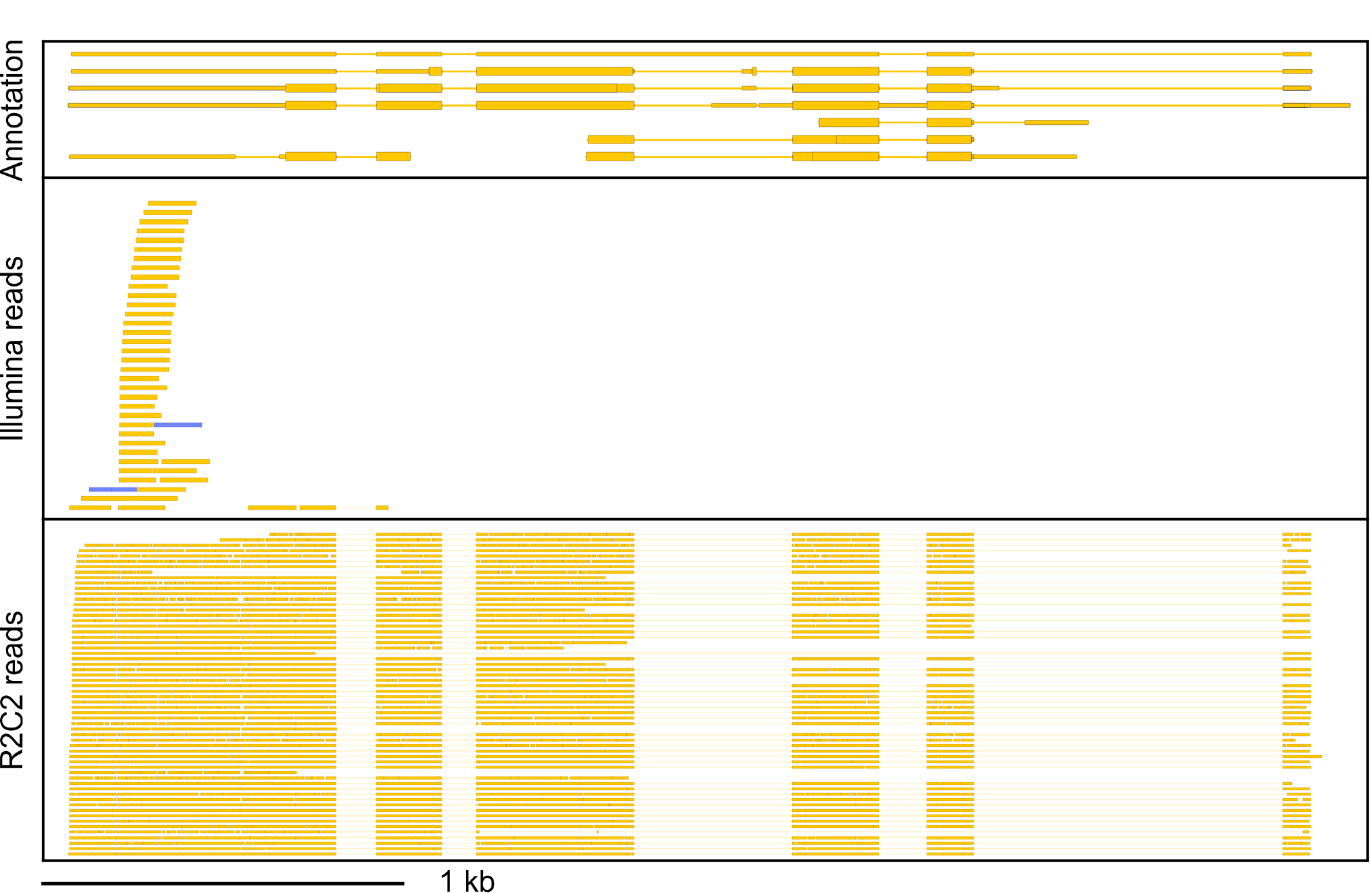
R2C2 reads sequence 10X full-length cDNA transcripts. Genome Browser shots of ACTB. Genome annotation is shown on top and Illumina reads (center) R2C2 reads (bottom) aligning to the locus are shown below. Both Illumina and R2C2 read alignments were randomly subsampled to 60 reads. The directionality of features is indicated by color (“top strand”=blue, “bottom strand”=yellow). Data for replicate 1 are shown.

**Table S1:** Oxford Nanopore Technologies sequencing run and read numbers. Values in parentheses indicate the percentage of raw reads being successfully converted into consensus reads. Note that R2C2 Consensus read numbers indicate consensus reads prior to post-processing. R2C2 Consensus reads after post-processing are given in Table 1.

| **Replicate 1** | **Basecalled raw reads** | **Median raw read len** | **R2C2 Consensus reads** |
| --- | --- | --- | --- |
| PromethION run 1 | 14,321,713 | 3601 | 8,588,694 (59.97%) |
| MinION run 1 | 2,476,880 | 4515 | 2,143,624 (86.55%) |
| PromethION run 2 | 12,069,611 | 3767 | 7,140,994 (59.17%) |
| MinION run 2 | 660,975 | 4074 | 547,384 (82.81%) |
| **Total** | **29,529,179** |  | **18,420,696 (62.38%)** |

**Table S1: Oxford Nanopore Technologies sequencing run and read numbers.**
| **Replicate 2** | **Basecalled raw reads** | **Median raw read len** | **R2C2 Consensus reads** |
| --- | --- | --- | --- |
| PromethION run 1 | 21,660,888 | 2746 | 13,603,254 (62.80%) |
| MinION run 1 | 4,865,719 | 2980 | 3,520,670 (72.36%) |
| **Total** | **26,526,607** |  | **17,123,924 (64.55%)** |

**Table S2:**
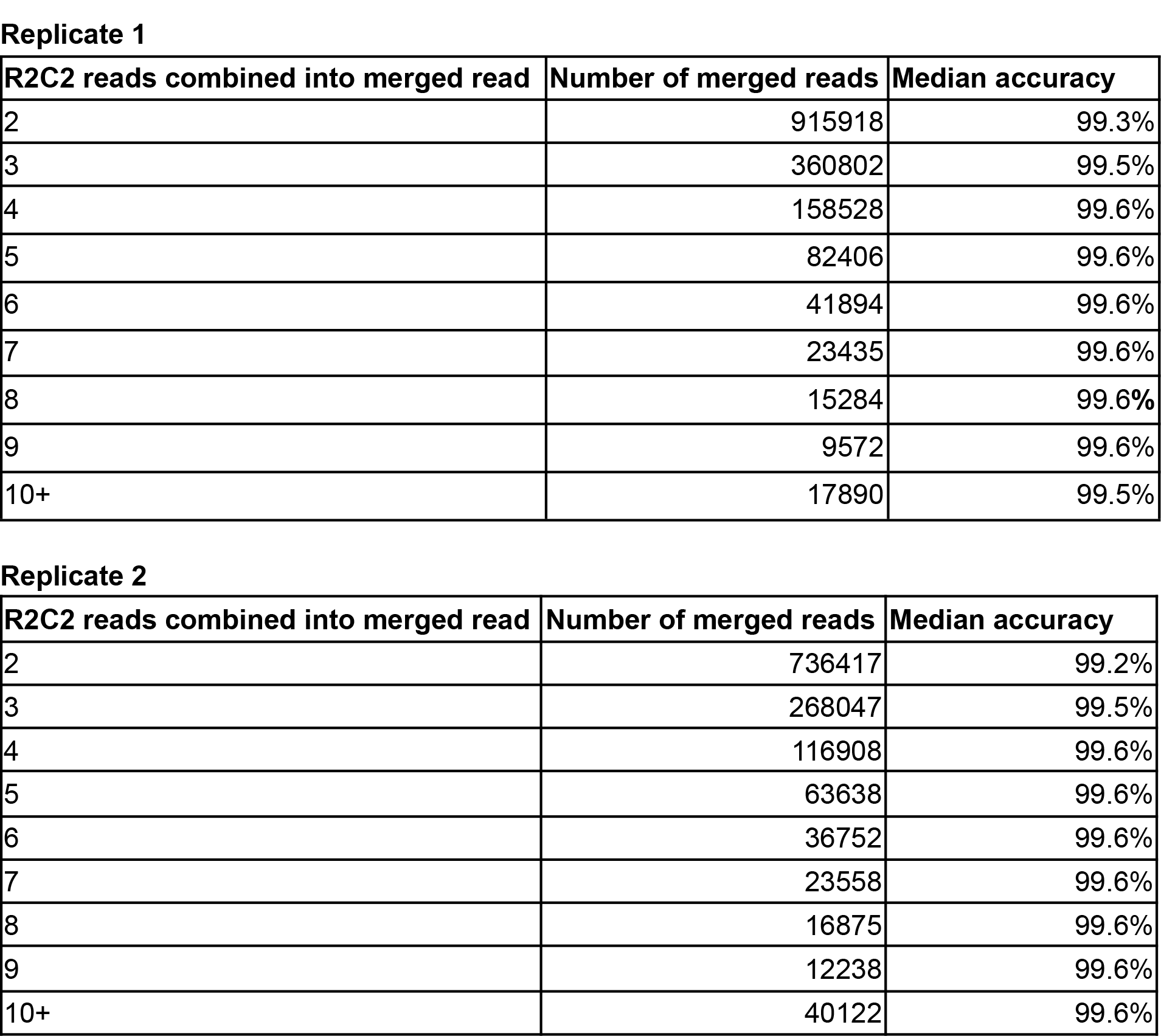
UMIs allow the merging of R2C2 reads originating from the same cDNA molecule.

**Replicate 1**
| **R2C2 reads combined into merged read** | **Number of merged reads** | **Median accuracy** |
| --- | --- | --- |
| 2 | 915918 | 99.3% |
| 3 | 360802 | 99.5% |
| 4 | 158528 | 99.6% |
| 5 | 82406 | 99.6% |
| 6 | 41894 | 99.6% |
| 7 | 23435 | 99.6% |
| 8 | 15284 | 99.6% |
| 9 | 9572 | 99.6% |
| 10+ | 17890 | 99.5% |

| **R2C2 reads combined into merged read** | **Number of merged reads** | **Median accuracy** |
| --- | --- | --- |
| 2 | 736417 | 99.2% |
| 3 | 268047 | 99.5% |
| 4 | 116908 | 99.6% |
| 5 | 63638 | 99.6% |
| 6 | 36752 | 99.6% |
| 7 | 23558 | 99.6% |
| 8 | 16875 | 99.6% |
| 9 | 12238 | 99.6% |
| 10+ | 40122 | 99.6% |

